# Ecological memory of prior nutrient exposure in the human gut microbiome

**DOI:** 10.1101/2021.06.21.448853

**Authors:** Jeffrey Letourneau, Zachary C Holmes, Eric P Dallow, Heather K Durand, Sharon Jiang, Savita K Gupta, Adam C Mincey, Michael J Muehlbauer, James R Bain, Lawrence A David

## Abstract

Many ecosystems retain an ecological memory of past conditions that affects responses to future stimuli. However, it remains unknown what mechanisms and dynamics may govern such a memory in microbial communities. Here, in both a human dietary intervention cohort and an artificial gut, we show that the human gut microbiome encodes a memory of past carbohydrate exposures. Changes in the relative abundance of primary degraders were sufficient to enhance metabolism, and these changes were mediated by transcriptional changes within hours of initial exposure. We further found that ecological memory of one carbohydrate species impacted metabolism of others. These findings demonstrate that the human gut microbiome’s metabolic potential reflects dietary exposures over preceding days and changes within hours of exposure to a novel nutrient.

**One Sentence Summary:** Recent nutrient exposures are encoded into the structure and activity of human gut microbial communities, which enables more efficient future metabolic responses.

## Main Text

Ecological memory describes a broad range of phenomena in which past disturbances experienced by an ecosystem influence community responses in the present (*1*). Such responses may reflect prior abiotic experiences (e.g. past rain or fires shaping reproduction strategies in plants) or biotic ones (e.g. species extinction altering competitive phenotypes among extant organisms) (*2*). Knowing that ecosystems retain memory has helped shape frameworks for reintroducing locally extinct species and promoting species diversity in settings like hardwood forests (*2, 3*). Ecological memory may also amplify the severity of ecosystem damage caused by climate change (*4*).

Despite the power of ecological memory to understand and manage ecological dynamics, our knowledge of how this concept applies to microbial ecosystems remains sparse. Within individual microbial species, evidence exists for memory-like processes. Among bacterial monocultures, past environmental conditions like nutrient availability affects metabolic potential (*5–7*), or the ability to utilize a substrate of interest. Yet, at the microbial community level, our understanding of how past environmental conditions affect future ecosystem function is limited. It has been observed that lasting changes in the abundance of taxa result from disturbances like oil spills (*8, 9*), antibiotic administration (*10*), dietary oscillations (*11*), and infection (*12*). In the human gut microbiome, it has been demonstrated that infection^13^ and obesogenic diet exposure^12^ induce a memory that affects the ecological outcome of subsequent perturbations. Still, it is not yet known what independent role microbes play in this memory (i.e. in the absence of host factors), nor the mechanism and properties of this memory. Furthermore, the fastest time scales on which microbial communities form ecological memory remain undefined. Due to the reproductive rates of bacteria, ecological memory may form in microbial ecosystems orders of magnitude faster than in communities of plants and animals (*13*).

Nutrient metabolism in the human gut provides an opportunity to test for ecological memory in a microbial community setting. Competition for nutrients in the gut is among the strongest microbial ecological forces (*14*) and host intake of nutrients varies on a daily basis (*15*). We thus reasoned that if the metabolic potential of the human gut microbiome reflects recent nutritional availability, newly introduced nutrients would be incompletely metabolized and require multiple exposures to be fully utilized by intestinal bacteria. To test this hypothesis, we enrolled 40 participants in a randomized placebo-controlled dietary intervention study. Individuals in the Prebiotic group were fed 18 g/day of inulin (Fig. S1a-d), a nutrient that can be metabolized by the gut microbiome (*16–18*), but which is typically consumed by individuals at low amounts (1-4 g/day) (*19*). We measured metabolic potential before and after inulin exposure using an *ex vivo* assay of the capacity for the fecal microbiome to degrade carbohydrates over the course of 24 hours (*20*). We observed a significant increase in inulin degradation after participants in the prebiotic group consumed inulin for one day (p = 0.0282, mixed-effects GLM; Fig. 1a, Fig. S2a). This finding suggests that within 24 hours of exposure to a novel nutrient, the human gut microbiome enhances its metabolic potential for that nutrient.

**Figure 1.**
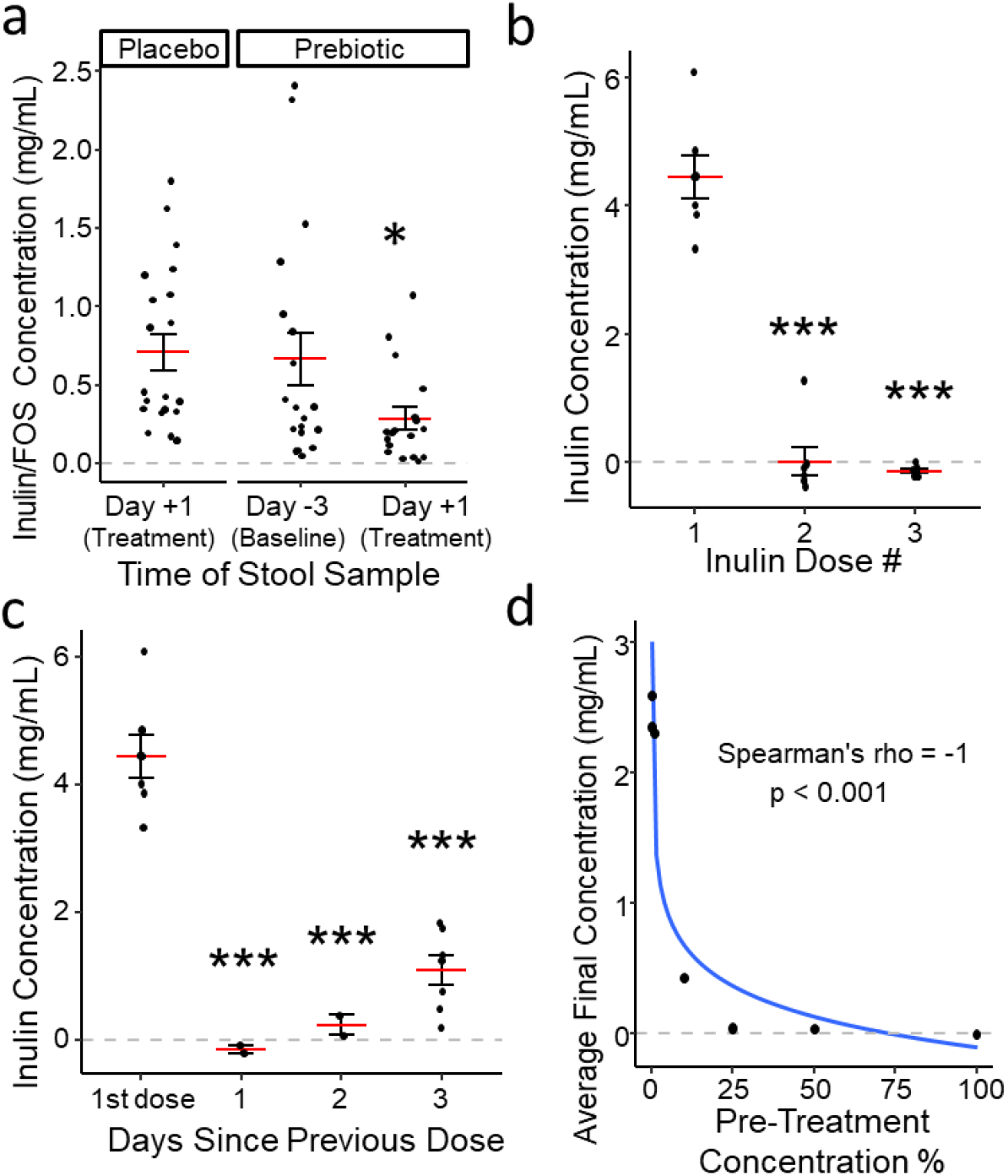
Ecological memory of prior nutrient exposure is encoded within 24 hours. **a**, Inulin/FOS concentrations after incubation of slurried stool samples with inulin for 24 hours *ex vivo*. (p = 0.0282, mixed-effects GLM; n = 21 participants in Placebo group, 19 in Prebiotic group Baseline, 18 in Prebiotic group Treatment.) **b**, Concentration of inulin remaining in each artificial gut vessel 6 hours after dosing. (Mixed-effects linear model with Dose 1 as intercept; n = 7 artificial gut vessels.) **c**, Concentration of inulin remaining in each vessel after 6 hours by time since previous dose. (Mixed-effects linear model with 1^st^ dose as intercept; n = 7, 2, 2, and 7 vessels, respectively. Setting the model intercept to 1 day shows a significant difference (p = 0.0475) between 1 and 3 days.) **d**, Average final inulin concentration (of triplicate cultures) after 6 hours incubation on inulin, preceded by pre-treatment with an inulin dose of varying concentration. Log2 regression line plotted. **a-c,** Mean and standard error plotted. * p < 0.05, ** p < 0.01, *** p < 0.001.

To systematically examine how inulin exposure altered the metabolic potential of the human gut microbiome, we considered three primary facets of ecological memory: lag, duration, and strength (*1*). Lag refers to the amount of time after a disturbance before the event is translated into a differential response to future stimuli. To measure memory lag with greater resolution and control than possible among humans *in vivo*, we employed an “artificial gut” model of the distal colon seeded with a fecal microbiome sample from a healthy human donor (Fig. S1e-f) (*21, 22*). To measure lag, we dosed each artificial gut vessel with a single 2 g dose of inulin each day (Fig. S1e). We measured differential responses to inulin supplementation by direct measurement of inulin concentration at −2 hr and +6 hr from each dose, as well as by continuous monitoring of pH, which decreases as inulin is metabolized (*23*) and impacts the overall ecology of the gut microbiome (*24*). These measurements revealed that by the second day of inulin treatment, gut microbial metabolism was altered; significantly less inulin remained in vessels at +6 hr on the second and third days of dosing (p < 0.001, mixed-effects linear model, Fig. 1b). Likewise, ecosystem pH reached a significantly lower minimum value on the second and subsequent days of dosing compared to the first (p < 0.001, mixed-effects linear model, Fig. S2b). Together, these findings suggest the gut microbiome can encode memory to a nutritional stimulus within a day of exposure.

We next investigated the duration and strength of microbiome memory. We added a second dosing week to our artificial gut model in which we varied the length between doses (Fig. S1e). Extending this period to two days between doses reduced, but did not entirely negate, the enhanced pH response (Fig. S2c). Microbiome potential to degrade inulin persisted even longer and remained enhanced when doses were separated by three days (p < 0.001, mixed-effects linear model; Fig. 1c). Still, our model suggested that inulin degradation capability had significantly decreased at 3 days between doses (p = 0.0175). We then set up a high-throughput *in vitro* anaerobic batch culture system to measure how the strength of microbiome memory varies following a wide range of inulin exposures (*25*). We observed a negative dose-dependence between concentration of inulin pre-treatment and subsequent metabolic memory (Spearman correlation p < 0.001; Fig. 1d, Fig. S2d). Notably, maximal inulin breakdown efficiency and acidification were reached below our original dose, which suggested that the gut microbiome’s ability to be primed for inulin metabolism could be saturated.

Having observed changes in community-level metabolism, we employed a multi-omics approach to determine the extent to which repeated nutrient exposure impacted community composition and function. While none of the taxa analyzed by 16S rRNA sequencing were found to have changed in abundance following the first inulin exposure (Dose 1 Hr +6), forty-three (47%) had altered abundance following the second exposure (95% credible interval excluding zero, Bayesian multinomial logistic-normal model; Fig. 2a, Fig. S3). Similarly, RNA sequencing revealed that across the global transcriptome (i.e. the collective transcriptome of the whole community), only two gene groups, both glycoside hydrolases, were differentially expressed after the first inulin dose; following the second dose, 18 genes were differentially expressed (p < 0.05, ALDEx2 GLM with Benjamini-Hochberg correction; Fig. 2b). Analyzing transcriptional changes within taxa, we found that the initial set of differentially expressed transcripts was significantly enriched for carbohydrate metabolism and transport functions (COG category “G”; p = 0.00779, chi-squared test; Fig. S4g). Following Dose 2, there was a 2.6-fold change in the number of transcripts with expression changes (p < 0.05, ALDEx2 GLM with Benjamini-Hochberg correction), and an additional 19 taxa (a 50% increase) had at least one such transcript (Fig. S4d).

**Figure 2:**
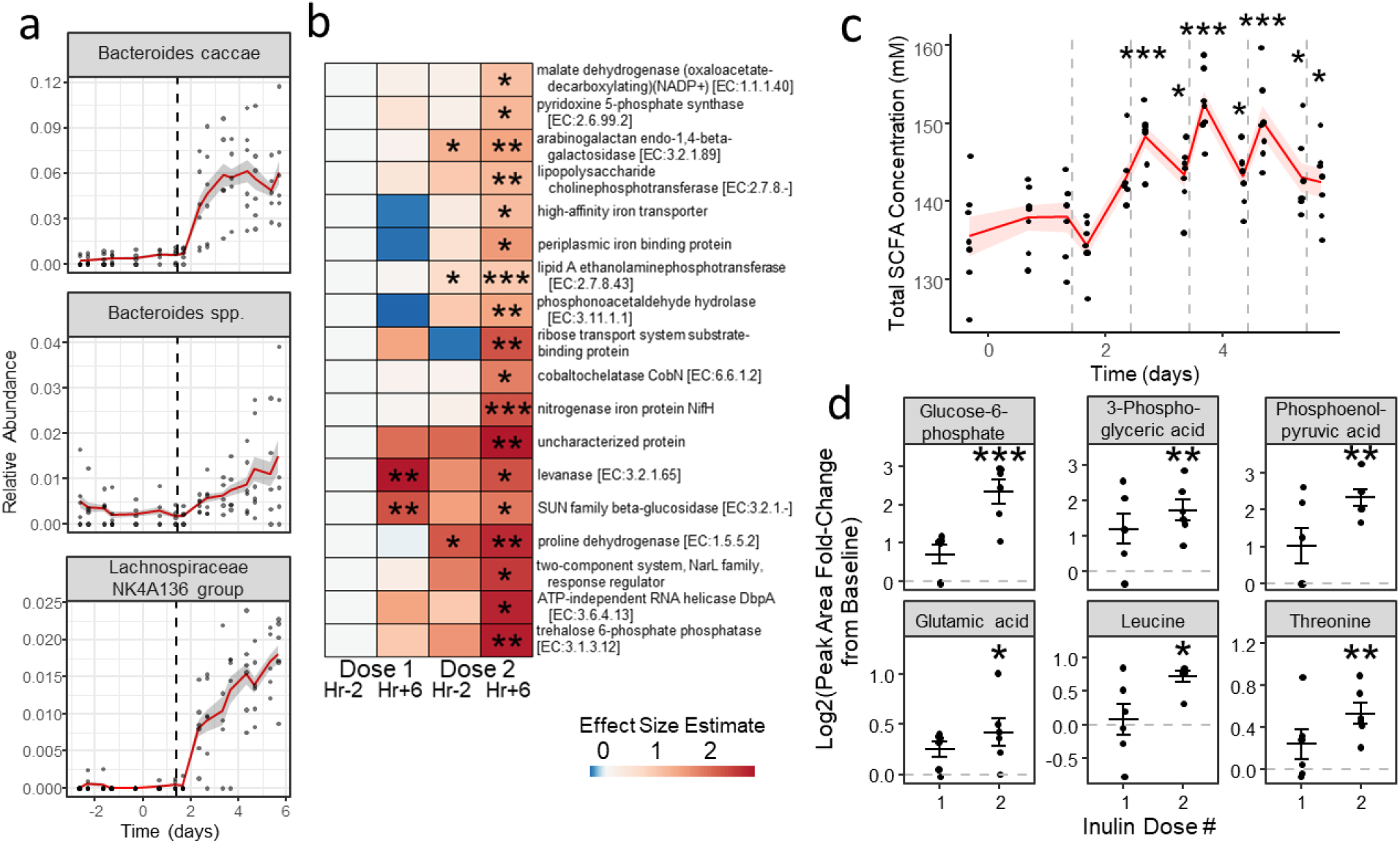
Compositional and transcriptomic changes reflect nutritional memory. **a**, Select SVs found to be significantly increased with inulin treatment by16S rDNA sequencing. (Multinomial logistic-normal model; n = 6 vessels.) **b**, Heatmap of KEGG gene functions found to be differentially expressed by time point in the global transcriptome. Color scale denotes effect size from baseline value within each KEGG function. (ALDEx2 GLM significance from baseline shown; n = 5 vessels.) **c**, Total SCFA concentration in artificial gut vessels over time. Vertical lines represent times of inulin dosing. (Mixed-effect linear model with Dose 1 Hr −2 (Day 1.42 on x-axis) as intercept; n = 7 vessels.) **d**, Change in concentration of select metabolites relative to baseline. Top row: fructose metabolism intermediates. Bottom row: amino acids. (Mixed-effects linear model significance from baseline shown; n = 6 vessels.) Mean and standard error plotted. * p < 0.05, ** p < 0.01, *** p < 0.001.

Metabolomic analyses further confirmed widespread alterations to microbial biochemical activity and environment following repeated nutrient exposure. Total short-chain fatty acid (SCFA) content was not significantly altered after the initial inulin dose, but was significantly increased after the second (p < 0.001, mixed-effects linear model; Fig. 2c), a trend which was driven by increases in acetate and butyrate (Fig. S5a). Since SCFA production is a major driver of acidification in the gut (*26*), these observations may explain the observed lag in pH decrease (Fig. S2b-d). Untargeted metabolomics revealed a 15.5-fold increase (2 to 31) in the number of metabolites whose levels changed after the second dose of inulin relative to the first (p < 0.05, mixed-effects linear model; Fig. 2d, Fig. S5b). Notably, the set of metabolites that increased after the second dose included fructose breakdown intermediates and five amino acids (p < 0.05, mixed-effect linear model; Fig. 2d, Fig. S5b), which suggests a shift from proteolytic metabolism towards a more saccharolytic state (*27, 28*). More broadly, our multi-omic analyses point to significantly greater changes in microbiome composition and activity as being associated with repeated nutrient exposure.

We investigated the specific ecological shifts that could amplify microbiome responses to a second inulin dose. We did not observe evidence that extracellular secretions induced memory, as inulin metabolism among inulin-naïve cultures could not be enhanced by adding conditioned media from inulin-treated cultures (Fig. 3a). We also did not find that changes to overall cell density in culture was related to inulin metabolic rates (Fig. S6a-b). By contrast, we were able to positively associate community taxonomic changes with microbiome memory to inulin exposure. Our 16S rRNA analysis of artificial gut communities in the two hours prior to the second inulin exposure revealed that 18 out of 92 analyzed SVs were significantly altered in abundance (95% credible interval excluding zero, Bayesian multinomial logistic-normal model; Fig. S3) relative to two hours prior to the first dose. These SVs included *Bacteroides caccae*, a known primary degrader of inulin (Virtual Metabolic Human (VMH)) (*29*), as well as *Bacteroides spp*. and *Bifidobacterium spp*., two genera that contain inulin-degrading species, and taxa such as *Lachnospiraceae NK4A136 group previously shown to be associated with intestinal SCFA levels* (*30*) *(Fig. 2a)*. Adding a *B. caccae* isolate derived from our artificial gut community to inulin-naïve stool-derived mixed community cultures was sufficient to enhance inulin metabolism (Fig. 2b), supporting the hypothesis that changes to taxonomic composition alone may drive memory in microbial communities.

**Figure 3:**
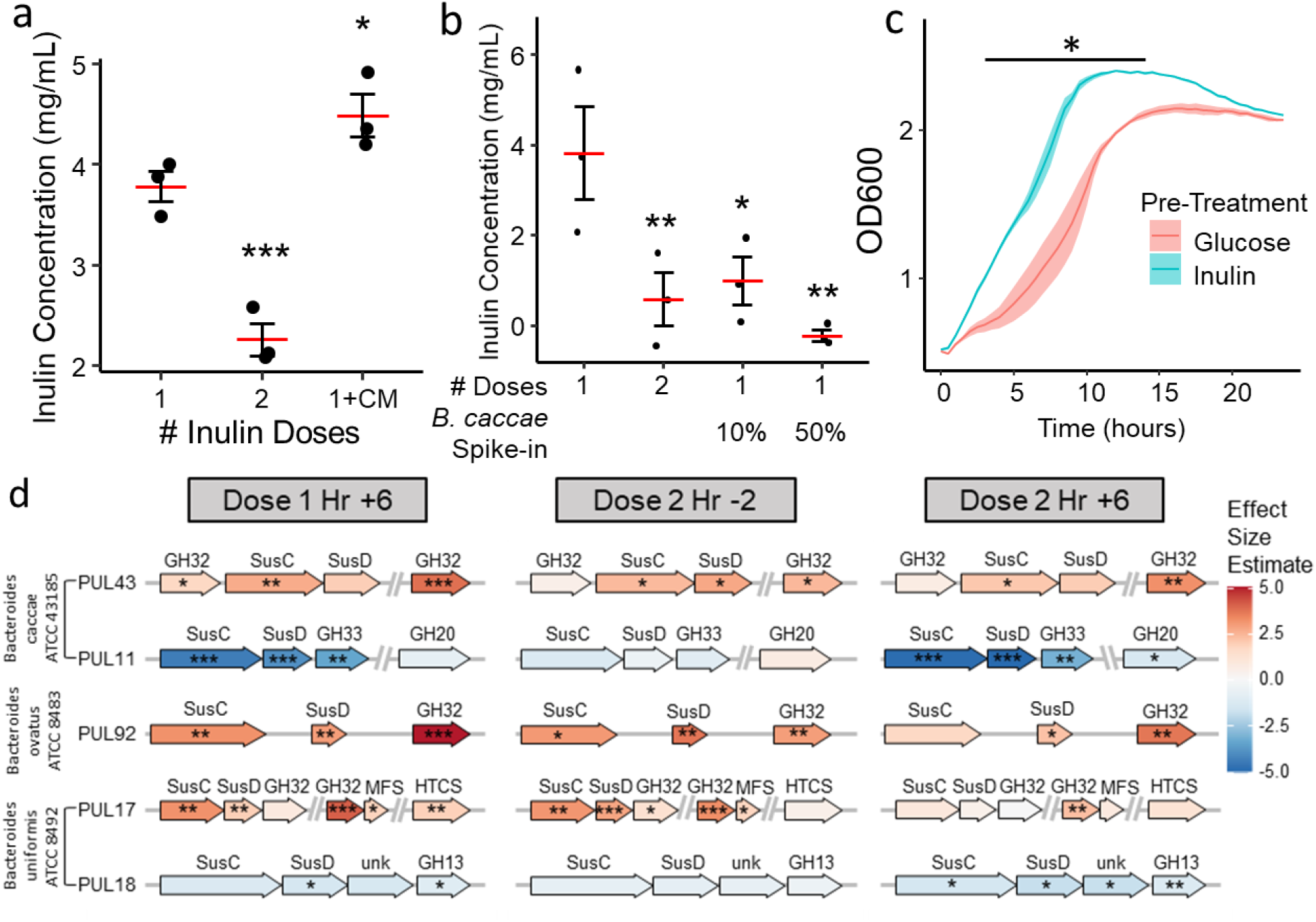
*Bacteroides* PUL activation precedes broader transcriptional and compositional changes. **a**, Conditioned media from inulin-exposed cultures did not increase metabolism of inulin in treatment-naïve cultures. (Linear model; n = 3 cultures). **b**, Effects of 10 or 50% *B. caccae* addition to complex community cultures. (Linear model; n = 3 cultures). **c**, Effects of glucose or inulin pre-treatment on growth curves of *B. caccae* monocultures grown on inulin in conditioned media (from inulin-naïve cultures). (Linear model; interaction term of time and pre-treatment p < 0.05 for all points below the line; n = 3 cultures). **a-c**, Mean and standard error shown. **d**, Activation and repression of select PULs in *Bacteroides* by time point. (ALDEx2 GLM; n = 5 artificial gut vessels). NS p > 0.05, * p < 0.05, ** p < 0.01, *** p < 0.001.

Given the known role of polysaccharide utilization loci (*31*) (PULs) in carbohydrate sensing among individual gut microbes (*32*), we also expected to observe changes among select microbes after initial exposure to inulin in our artificial gut. Indeed, we found activation/repression of PULs in twelve *Bacteroides* taxa in the hours preceding the second inulin dose (Fig. S7). Since our analysis normalized expression levels within taxa, these differences reflected changes in the expression of genes within a given genome and not simply changes in bacterial taxonomic abundance. These PULs included genes encoding inulin-degradative glycoside hydrolase family 32 (GH32) enzymes, which were also upregulated in the global transcriptome after a second inulin dose (Fig. 2b), as well as transcriptionally linked SusC/SusD homologs, which work together to bind and import oligosaccharides (*32*) (Fig. 3d, Fig. S7). Given, however, that these PULs were activated even earlier, at Dose 1 Hr +6 (Fig. 3d, Fig. S7), we suspected that transcriptional changes played a role in enabling growth on the new substrate. In support of this hypothesis, we found that even after controlling for starting cell density, inulin pre-treatment enhanced the ability of *B. caccae* to grow on inulin (Fig. 3c, Fig. S6c). We therefore suspect that patterns of transcriptional change observed when monocultures of bacteria leave lag phase during diauxic shifts (*33*) are likely to also occur among *Bacteroides* species in a mixed species setting.

We also identified upregulated sets of syntenic genes outside of the *Bacteroides*, identifying several putative carbohydrate-associated loci (p < 0.05, ALDEx2 GLM with Benjamini-Hochberg correction; Fig. S4e) in taxa that degrade products of inulin hydrolysis (glucose, fructose, sucrose, or short-chain fructo-oligosaccharides (scFOS)) (*29*). Seven *Bacteroides* and two *Clostridium* species had loci with at least one gene still significantly upregulated two hours prior to the second prebiotic dose (Figs. S4e, 6d). Three of these *Bacteroides* taxa maintained activation of genes encoding inulin degrading enzymes, and *B. caccae* also continued to upregulate a fructokinase gene (p < 0.05, ALDEx2 GLM with Benjamini-Hochberg correction; Fig. S6d). This genetic activity, coupled with enhanced degradation of inulin (Fig. 1b), may have enabled additional taxa to upregulate genes involved in pathways downstream of inulin processing (*34*) (Fig. 3b) and increase SCFA production (Fig. 2c, Fig. S5a). Our metagenomic and metabolomic data suggest that sustained expression of GH32 enzymes, along with increased relative abundance of the taxa that produce them (Fig. 2a), prime microbial communities for a more widespread metabolic response to inulin upon re-exposure.

Given that expression of a particular PUL or growth of a certain taxon may be activated by multiple substrates (*29, 35*), we reasoned that ecological memory of inulin may trigger, or be triggered by, ecological memory of related carbohydrate compounds. We found evidence for this model in our human dietary intervention study. Participants’ total baseline dietary fiber intake, as estimated by Diet History Questionnaire III (DHQ3), was negatively correlated with fecal inulin/FOS content in donor stool one day after the start of treatment in the Prebiotic group (p = 0.0232, Spearman correlation; Fig. 4a) but not the Placebo group (p = 0.402, Spearman correlation; Fig. S8d). Thus, participants who naturally consumed a high rate of dietary fiber tended to excrete less undigested inulin than individuals with low habitual fiber consumption.

**Figure 4:**
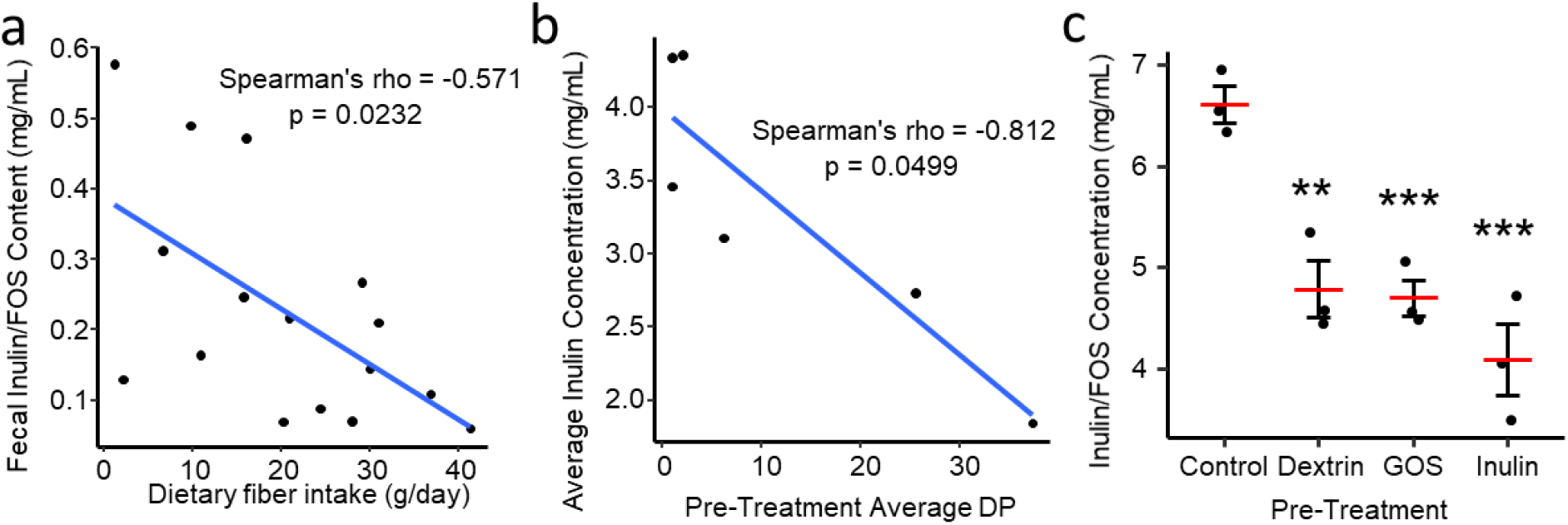
Ecological memory is cross-reactive. **a**, Correlation between participant baseline dietary fiber (DHQ3) intake and fecal inulin/FOS content on Treatment Day +1 in the Prebiotic group. (n = 16 participants.) **b**, Correlation between average DP of pre-treatment inulin or inulin constituent and average final inulin concentration (or triplicate cultures). **c**, Final inulin concentration after pre-treatment with different prebiotics. (Linear model with control as intercept; n = 3 cultures.) Mean and standard error plotted. * p < 0.05, ** p < 0.01, *** p < 0.001.

Next, we tested for the cross-reactivity of ecological memory to inulin degradation using *in vitro* models. We treated human gut microbiome cultures with the constituent components of inulin (fructose, glucose, sucrose, and FOS/inulin of various chain lengths). This treatment indeed enhanced subsequent inulin metabolism to varying degrees (p < 0.05, linear model; Fig. S8b-c). The magnitude of this effect was correlated with component chain length (Spearman correlation, p = 0.0499; Fig. 4b). We next tested whether treatment with non-fructan polysaccharides could enhance inulin metabolic rates. We found exposure to galacto-oligosaccharides (GOS) and dextrin resulted in partial, significant increases in inulin degradation capacity (p < 0.01, linear model; Fig. 4c) and significant decreases in culture pH relative to control following inulin treatment (p < 0.001, linear model; Fig. S8a). Community analysis of microbiome cultures revealed that non-inulin treatment effects on the abundance of a specific bacterial taxon, *Clostridium ramosum*, were correlated with subsequent inulin response (p < 0.0130, Spearman correlation; Fig. S9a). *C. ramosum* is a known degrader of scFOS, as well as the component mono- and di-saccharides of GOS and dextrin, respectively (*29*). This correlation therefore supports a model where ecological memory within microbiome can exhibit cross-reactive properties in which altering the abundance of a nutrient-degrading microbial species ultimately impacts the rates at which related nutrients will subsequently be metabolized.

In concert, our findings demonstrate the existence of ecological memory of past nutrient exposure in the human gut microbiome. Previously, functional changes in the gut microbiome have generally been thought to take days to weeks to change (*16*), with evidence of transcriptomic changes within an individual’s gut microbiome on the order of days (*36*). Our findings suggest ecological memory can be encoded even more rapidly, on sub-daily time scales for the human gut microbiome. This response time is consistent with how quickly individual microbes are known to undergo diauxic shifts, which also occur on the order of hours (*37*). Indeed, it is likely that selective pressures for rapid microbial metabolic change exist in environments like the mammalian gut, where community members replicate on hourly timescales and nutrient availability varies in both stochastic and rhythmic manners within a single day (*38*). Such adaptation may also benefit hosts by providing them with an adaptative microbial response to dietary shifts (*36*). Yet, given the benefits to rapid metabolic plasticity, it is perhaps surprising that we also observe evidence that after a nutrient is withdrawn, ecological memory persists for days and therefore likely across multiple generations of bacteria. Bacterial communities or their members may benefit from “bet hedging” strategies that balance the odds that a withdrawn nutrient is reintroduced (*33*). It is also possible that once the transcriptional, compositional, and metabolomic landscape of a microbial community has been altered, restoring to an original starting state will encounter delay (*39*).

Our finding of ecological memory in human gut microbial systems also suggests avenues for the rational design of treatments that alter how the gut microbiome harvests energy (*40*) or metabolizes drugs (*41, 42*). To date, individualized therapies have accounted for inter-individual variation in microbiome composition and function (*43*), which have been linked to fixed population differences in both overall diversity and specific taxonomic differences (*44*). Yet, if the metabolic potential of the gut microbiome is plastic, observed microbiome heterogeneity may also reflect recent *intra*-individual variation in behavior or lifestyle. Rationally designed therapies may therefore benefit from monitoring changes to, and even manipulating, microbiome metabolic potential over time (*45*). For example, our work suggests that microbiome-targeting nutritional interventions have the most potential to impact the microbial metabolism of individuals who are normally deficient in that nutrient’s intake (Fig. 4a). Moreover, our finding that microbiome memory does not persist indefinitely suggests that such interventions will require repeated administration to sustain their microbial ecological impact.

## Acknowledgments

We would like to thank Justin Silverman for assistance with sequencing and model design; Andrew Grover for help sampling from the artificial guts; Verónica Palacios for work on the human research portion of the study; Danielle Anderson and Ken Racicot for providing the snack bars used in the study; Neil Surana for helpful guidance related to our literature review; Tonya Snipes, Lisa Alston-Latta, and Margaret Huggins for keeping our lab spaces and glassware clean; and our study volunteers for their participation.

## Funding

This work was supported by:

National Institutes of Health grant 1R01DK116187

Office of Naval Research grant N00014-18-1-2616

Translational Research Institute through Cooperative Agreement NNX16AO69A

Damon Runyon Cancer Research Foundation

University of North Carolina CGIBD (NIDDK P30DK034987)

This study used a high-performance computing facility partially supported by grant 2016-IDG-1013 (HARDAC+: Reproducible HPC for Next-Generation Genomics) from the North Carolina Biotechnology Center.

## Author contributions

Conceptualization: JL, ZCH, JRB, LAD

Data curation: JL, MJM

Formal analysis: JL

Funding acquisition: LAD

Investigation: JL, ZCH, EPD, HKD, SJ, SKG, ACM, MJM

Software: JL

Visualization: JL

Writing – original draft: JL

Writing – review & editing: JL, LAD

## Data and materials availability

Sequencing data are publicly available via the European Nucleotide Archive with the accession numbers PRJEB45244 (metatranscriptomics) and PRJEB45247 (16S sequencing). Additional data and code are available upon request from the corresponding author.

## Supplementary Materials

Materials and Methods

Figs. S1 to S10

References (*46–58*)

