## Supplemental Methods and Figures for "Ecological memory of prior nutrient exposure in the human gut microbiome"

**This PDF file includes:**

Materials and Methods
Figs. S1 to S10
References (46-58)

### Materials and Methods

Carbohydrate sources

The following sources of fructan prebiotics were used in this study: Synergy1 inulin (Orafti), inulin from dahlia tubers (Sigma), fructo-oligosaccharides from chicory (Sigma), inulin (NOW), and inulin FOS (Jarrow) (Extended Data Fig. 10). Orafti Synergy1 inulin was used for the human study as a food-safe ingredient that could be added to the snack bars, Sigma inulin from dahlia tubers was used for *in vitro* work including the artificial gut run as a relatively pure long-chain inulin. Jarrow inulin FOS was used for its greater solubility in select *in vitro* experiments (Figs. 3c, 4c, Extended Data Figs. 6a-c, 9a), where it was necessary to measure OD600 and/or ensure that pre-treatment inulin could be removed (via supernatant) prior to the treatment phase.

The following sources of non-fructan prebiotics were used: dextrin (Benefiber) and galacto-oligosaccharides (Bimuno). The following simple sugars were also used: dextrose (Amresco), fructose (Amresco), and sucrose (Sigma).

Participant recruitment and sample collection

We recruited 40 healthy subjects by use of flyers on Duke University campus as well as electronic postings on DukeList (a university-internal classifieds website), a lab website, and ClinicalTrials.gov. The study protocol was approved by the Duke Health Institutional Review Board (IRB) at Duke University under protocol number Pro00093322. All patients provided written informed consent via an electronic consent form (eConsent) prior to participation in the study protocol. The study protocol was registered on ClinicalTrials.gov, with the identified number NCT04055246.

All participants were between the ages of 18 and 35, weighed at least 110 pounds (50 kg), and had a body mass index (BMI) between 17 and 27.5. Furthermore, individuals were excluded from participation if they: had a diagnosis of psychiatric or neurological disorder, were currently on steroid medications, had used recreational drugs within the past month, consumed at least 2 alcoholic beverages per day on average, had dietary restrictions of milk or dairy products, had food allergies to wheat/gluten/nuts/soy, or were currently pregnant or breastfeeding. A secondary screen further excluded individuals who scored greater than 9 on a modified Patient Health Questionnaire-9 (PHQ-9, a depression severity questionnaire), had a colonoscopy or oral antibiotics within the past month, or had a history or current diagnosis of any of the following: irritable bowel syndrome, inflammatory bowel disease, type 2 diabetes, chronic kidney disease with decreased kidney function, intestinal obstruction, or untreated colorectal cancer.

Participants were 60% female (24/40) and 40% male (16/40). Participants were 57.5% white (23/40), 40% Asian (16/40), 2.5% Black (1/40), 2.5% Native American (1/40), 2.5% Native Hawaiian or Other Pacific Islander (1/40), and 12.5% Hispanic or Latino (5/40). Average age at time of enrollment was 25.6 ± 4.9 years (mean ± stdev). Most participants (67.5%; 27/40) were omnivores, 17.5% were vegetarian (7/40), 12.5% ate everything except red meat (5/40),

2.5% did not respond to this question (1/40), and none were vegan. Average weight was 147.9 ± 21.6 lb and average BMI was 23.2 ± 2.2.

As shown in Extended Data Fig. 1a, participants provided stool samples twice weekly on Tuesday and Friday over a three week study period. If a participant was unable to produce a sample on the requested day, they were instructed to provide the next available sample (on Wednesday or Thursday in place of a Tuesday sample, or on Saturday or Sunday in place of a Friday sample). Samples were self-collected using polypropylene scoop-cap tubes (Globe Scientific), and participants were instructed to keep samples in personal freezers until ready to transport to lab using a provided insulated container and ice pack. Once arrived, samples were kept at -20 °C for up to a week and then moved to -80 °C until further processing.

On the second study week, participants consumed treatment bars (Extended Data Fig. 1b) twice daily for five days (Monday through Friday) containing either 9 g/bar inulin (prebiotic group, n=19) or 9 g/bar maltodextrin (placebo group, n=21) (Extended Data Fig. 1d). Participants were blinded to which group they were in (Extended Data Fig. 1c). The bars were manufactured by the Natick Soldier Research Development and Engineering Center (NSRDEC) in Natick, MA, USA, based on a modified formula of the First Strike™ bar manufactured at NSRDEC.

All participants completed three dietary surveys. The Diet History Questionnaire III (DHQ3) was administered prior to the Baseline and assessed participants' eating habits over the past month. The Automated Self-Administered 24-Hour Dietary Assessment Tool (ASA24-2018) was administered twice, once during the Baseline week and once during the Treatment Week, to provide a log of everything participants ate in the day prior to taking the test.

In some instances, a participant did not provide a stool sample on the day requested or not enough sample was present for all analyses, and one participant did not complete the DHQ3. Thus, n is not always 19 for Placebo and 21 for Prebiotic; otherwise, no samples were intentionally excluded from analysis.

Measurement of metabolic potential in stool samples

To measure the capacity of microbial communities to degrade prebiotic substrate, we used a previously described stool fermentation assay (*20*). Briefly, stool samples were weighed out in an anaerobic chamber and combined with pre-reduced phosphate-buffered saline (PBS) at 10% weight/volume in polyethylene filter bags with 0.33-mm pore size (Whirl-Pak B01385). Samples were then homogenized (aerobically) using a stomacher (Seward Stomacher 80 MicroBiomaster) set to medium for 60 second. Total time exposed to oxygen did not exceed 10 minutes.

Either 1 mL of either PBS (control) or 1% (10 mg/ml) solution of Orafti Synergy1 inulin was added to 24-well plate wells. Then, 1 mL of stool slurry was aliquoted into each well (final concentration of inulin 5 mg/ml). Each stool sample was assayed in duplicate (2 control wells and 2 inulin wells). Plates were sealed with adhesive foil seals and incubated anaerobically for 24 hours at 37 °C. Aliquots were taken from each well and saved at -80 °C.

Artificial gut culturing and sampling

Artificial guts were run according to a previously established protocol (*22*). An eight-vessel continuous flow artificial gut system (Multifors 2, Infors) was used to culture gut microbes seeded from human stool samples. Replicate artificial gut vessels were uniformly inoculated with a starting community derived from a single healthy stool donor so that our analyses could assume measurement variation was due to technical sources of noise (*22*). Vessels were sterilized and prepared with 300 mL of fresh modified Gifu Anaerobic Medium (mGAM) (*46*). Nitrogen was sparged into the vessels at 1 L/min to maintain positive pressure and create an anaerobic environment. Vessels were inoculated using culture aliquots saved at -80 °C from a previous artificial gut run (original fecal samples obtained from a healthy volunteer who provided written informed consent per Duke Health IRB Pro00049498). From each of 12 frozen aliquots, 1.5 mL was added to 5 mL mGAM media and incubated for 6 hours at 37 °C in an anaerobic chamber (Coy). 5 mL of these cultures were then transferred to media bottles contain 100 mL mGAM and incubated overnight (17 hours) at 37 °C. After this final incubation, cultures were combined aseptically in a biosafety cabinet and loaded into syringes, with two 50 mL aliquots inoculated into each vessel via a septum in the vessel lid, for a final working volume of 400 mL. The media feed was started 24 hr after inoculation at a constant rate of 400 mL per day (a rough approximation of 24 hr average passage time in the human GI tract).

The pH of each vessel was monitored and controlled by the IRIS software (v6, Infors). Vessel pH was maintained within the ranges of 6.8 ± 0.1 during burn-in phase and 6.2 ± 0.7 thereafter using a 1 N HCl solution and a 1 N $H_3PO_4$ solution. pH was measured continuously with Hamilton EasyFerm Plus PH ARC 225 probes. The pH probes were calibrated with a two-point calibration with standardized pH buffers at 4.00 ± 0.1 and 10.00 ± 0.1 (BDH). Vessels were maintained at 37 °C via the Infors's onboard temperature control system. Vessels were continuously stirred at 100 rpm using magnetic impeller stir-shafts.

Prior to sampling, sampling lines were cleared with a 0.2 μm filter-tipped syringe and wiped clean with 70% ethanol. Sampling consisted of the collection of 7 mL of active artificial gut culture via sterile syringe and then immediate storage in labeled cryovials at -80 °C. Dosing was done by combining 16 g inulin with 80mL PBS in a biosafety cabinet, and administering 10 mL of the mixture to each vessel by needle-equipped syringe through a septum at the top of the vessel.

Samples from all seven inoculated vessels were analyzed for inulin content, pH, and SCFA content, as described below. Due to an issue with media flow on the final day of treatment (Day 11; Extended Data Fig. 1e), samples from this day were not analyzed for inulin content, and instead, samples from all 7 vessels on Day 8 were used to test the effect of 3 days between doses (Fig. 1c). Due to analytical pipeline throughput limitations, we prioritized samples from six of the vessels for metatranscriptomic sequencing and untargeted metabolomics. Additionally, DNA/RNA sequencing data from one of the artificial guts was excluded from statistical analyses involving the origins of ecological memory, due to this replicate exhibiting evidence for a delayed memory effect.

Small-batch bacterial culturing

To perform controlled *in vitro* experiments (Figs. 1d, 3a-c, 4b-c, Extended Data Figs. 2d, 6a-c, 8a-c, 9) in a more high-throughput way than the artificial gut allowed, we carried out small-batch anaerobic culture experiments based on previously established protocols (*25*). Cultures were started from a 1 mL frozen aliquot from a previous artificial gut run was added to 4 mL mGAM and grown anaerobically for 6-8 hours at 37 °C in a 14 mL polystyrene round-bottom tube. This 5 mL culture was then added to 45 mL mGAM and grown overnight (16-18 hours) at 37 °C. Following this overnight culture, culture tubes each containing 3 mL mGAM were inoculated with 1 mL overnight culture for a total volume of 4 mL. Cultures were pre-treated by adding 200 µL of carbohydrate at 100 mg/mL (except for the dose-response experiment), for a final concentration of 4.8 mg/mL. After 24 hours incubation, cells were spun down 10 min at 3000 x g and resuspended in fresh mGAM along with 200 µL of inulin at 100 mg/mL. Samples were collected after a final 6 hour incubation. Inulin concentration was analyzed by HPAEC-PAD as described below and endpoint pH was measured by Accument AR-15 pH meter (Fisher Scientific).

Conditioned media was generated by growing stool-derived microbiome cultures on mGAM overnight and filter-sterilizing the resultant culture. The *Bacteroides caccae* isolate used (strain ATCC 43185) was previously isolated from the community used for the artificial gut experiment. For the spike-in experiment (Fig. 3b), cultures were grown overnight, then diluted 1:10 and grown for 6 hours with or without inulin. OD600 was measured, and diluted with fresh mGAM to a combined calculated OD600 of 1.0 in each 4 mL culture, with *B. caccae* and whole community cultures mixed at 10/90 or 50/50% by OD600.

For the growth curve experiment (Fig. 3c, Extended Data Fig. 6c), cultures were pre-treated for 9 hours on (inulin-naïve) conditioned mGAM supplemented with 4.8 mg/mL glucose or inulin/FOS (Jarrow) at which point OD600 was measured and cultures were diluted to a calculated OD600 of 0.5 into 200 µL conditioned mGAM plus 10 µL of 100 mg/mL glucose of inulin (final concentration of 4.8 mg/mL) in a 96-well plate. OD600 was measured by SPECTROstar Nano Microplate Reader (BMG Labtech) in the anaerobic chamber, with 48 readings taken at 30 min intervals, with the plate shaken before each reading. A growth curve was fit using the growthcurver package in R (Extended Data Fig. 6c).

Quantification of inulin

Analysis of inulin content was performed by high-performance anion-exchange chromatography with pulsed amperometric detection (HPAEC-PAD). Briefly, 200µL of each sample (in randomized order) was added to 800µL of deionized (DI) water, ACS reagent grade, 18MΩ-cm resistivity. Samples were then centrifuged at 14,000 x g for 5 minutes at 4°C. The resulting supernatant was transferred to a 0.22 µm spin filter column and centrifuged at 14,000 x g for 5 minutes at 4°C. The filtrate was analyzed on a Dionex ICS-6000 HPIC system equipped with a Dionex AS-AP autosampler and a pulsed electrochemical detector consisting of an amperometric flow-through cell and a silver/silver chloride reference electrode. The electrochemical detector provided the following waveform: $E_1$ = +0.1V ($t_1$ = 0.00s - 0.40s), integration from 0.20 to 0.40 s, $E_2$ = -2.0V ($t_2$ = 0.41s – 0.42s), $E_3$ = +0.6V ($t_3$ = 0.43s), $E_4$ = -0.1V ($t_4$ = 0.44 – 0.50s).

Separation was carried out on a CarboPac PA200 analytical column equipped with a CarboPac PA200 guard column at 30°C. The autosampler was kept at 4°C to prevent degradation of carbohydrates. A gradient elution was performed using the following eluents: 100 mM NaOH (eluent A) and 1 M NaOAc, 100 mM NaOH (eluent B). The eluents were kept blanketed under nitrogen at 6psi to prevent the formation of carbonate. A flow rate of 0.5mL/min was used and the linear gradient was setup as follows: 98% eluent A and 2% eluent B from -5 to 15 min, ramping to 50% eluent A and 50% eluent B at 70 min, then ramping back to the original parameters at 70.1 min and remaining that way until the run was ended at 75 min.

The data was acquired and processed using Thermo Scientific Dionex Chromeleon Chormatography Data System software, using inulin (inulin from dahlia tubers, Sigma), glucose (Amresco), fructose (Amresco), sucrose (Sigma), 1-kestose (≥98%, Sigma), and nystose (≥98%, Sigma) standards to identify peaks. Resulting data was processed in R by transforming peak areas to concentrations based on the inulin standard for peaks with degree of polymerization (DP) >10, and based on an average coefficient derived from kestose and nystose standards for DP 3-10. Peak values for negative control samples containing no inulin were subtracted from bacterial culture samples. Negative controls were unavailable for stool samples due to the unique nature of each sample. Since Sigma dahlia inulin contained almost exclusively long-chain inulin (DP 11+; Extended Data Fig. 10b), we analyzed only this fraction for experiments using this inulin source. Otherwise, (i.e. for Orafti Synergy1 inulin and Jarrow inulin FOS), we included all inulin/FOS of DP3+ in our analysis.

16S rRNA sequencing

For all samples, 16S rRNA gene amplicon sequencing was performed using custom barcoded primers targeting the V4 region of the gene according to previously published protocols (20, *22*, *47*, *48*). Samples were randomized and DNA extractions were performed using the Qiagen DNeasy PowerSoil DNA extraction kit (ID 12888-100). Sequencing runs were standardized to 10 nM and sequenced using an Illumina MiniSeq with paired end 150 bp reads. DADA2 was used to identify and quantify sequence variants (SVs) in our dataset, using version 123 of the Silva database (*49*). We retained only samples with more than 5000 read counts to remove outlying samples that may have been subject to library preparation or sequencing artifacts, (*50*) and to only retain taxa that appeared more than three times in at least ten percent of samples.

Metatranscriptomics

RNA was extracted from samples using Quick-RNA Fungal/Bacterial Miniprep Kit (Zymo Research R2014) according to the manufacturer's instructions. Samples were then shipped to Novogene (Sacramento, CA) for further processing. mRNA was enriched using the Illumina Ribo-Zero Gold rRNA Removal Kit (Epidemiology) and 300 bp reads were sequenced on an Illumina HiSeq.

Analysis of sequencing reads was carried out using a pipeline adapted from previous metagenomic work (*13, 36*). Briefly, reads were matched using the Bowtie 2 sequence alignment program to a reference survey from version 3.5 of the Integrated Microbial Genomes system (*51*). We used the system's annotations to the COG (*52*) and KEGG (*53*), and EC (*54*) databases for functional analyses. For single-species level analyses, reads that mapped to more than 1 reference genome were discarded. For global analyses, however, reads that mapped to multiple reference genomes were still counted.

We initially detected RNA from 1394 taxa; this was reduced to 343 after retaining only taxa that appeared more than three times in at least ten percent of samples. The same filter was applied to genes within taxa; only genes that appeared more than three times in at least ten percent of samples were retained. Finally, we could only perform our analysis (ALDEx2 GLM, and PERMANOVA) on taxa that had at least two genes present (non-zero) in every sample. This filter brought the final number of taxa analyzed down to 178.

Metabolomic profiling

SCFAs were quantified by GC as previously described (*20*). Briefly, samples were acidified by adding 50 µL 6 N HCl to lower the pH below 3. The mixture was vortexed and then centrifuged at 14,000 x g for 5 min at 4 °C. 750 µLof this supernatant was passed through a 0.22 µm spin column filter. The resulting filtrate was then transferred to a glass autosampler vial (VWR, part 66009-882). Filtrates were analyzed on an Agilent 7890b gas chromatograph equipped with a flame ionization detector and an Agilent HP-FFAP free fatty-acid column (25 m x 0.2 mm [inner diameter] x 0.3 µm film). A volume of 0.5 µL of the filtrate was injected into a sampling port heated to 220 °C and equipped with a split injection liner. The column temperature was maintained at 120 °C for 1 min, ramped to 170 °C at a rate of 10 °C/min, and then maintained at 170 °C for 1 min. The helium carrier gas was run at a constant flow rate of 1 ml/min, giving an average velocity of 35 cm/s. After each sample, we ran a 1-min postrun at 220 °C and a carrier gas flow rate of 1 mL/min to clear any residual sample. All C2:C5short-chain fatty acids were identified and quantified in each sample by comparing to an 8-point standard curve.

Additional metabolites were quantified by untargeted GC/MS similar to previously described work (55) with some differences noted below. As above, samples were pre-processed by vortexing, centrifuging at 14,000 x g for 5 min at 4°C, and passing 750 µL of this supernatant through a 0.22 µm spin column filter. Proteins were then precipitated by adding 750 µL methanol (containing 6.25 mg/L C14:0-$D_{27}$ myristic acid as retention time standard) to 100 µL of each sample in a glass vial. After additional vortexing and centrifugation, 700 µL of each supernatant was transferred into a fresh glass vial for 4-6 hours of drying on a SpeedVac. Next, 100 µL toluene was added for a final step of azeotropic drying for 30 min, and the dried samples were stored at -80 °C for later derivatization. Subsequent steps included addition of 25 µL methoxyamine HCl with incubation at 50 °C for 30 min (for methoximation of reactive carbonyl groups) followed by addition of 75 µL N-methyl-N- (trimethylsilyl) trifluroacetamide (MSTFA, Cerilliant M-132) and similar incubation for trimethylsilyl replacement of exchangeable protons. Supernatants were transferred to GC vials and analyzed on a 7890B GC-5977B ei-MS (Agilent

Corp., Santa Clara, CA). We detected 547 metabolite features that were present in at least 10% of samples, from which 206 were annotated using our GC/MS spectral library (*55*, *56*) for use in subsequent statistical analysis (the remaining 341 included 280 unannotated, and 61 that either had uncertain annotation or were suspected contaminants). Levels for each metabolite were given as log-2 integrated peak area. For the purposes of statistical testing, we set the values of sample-metabolites that were missing, assumed below the limit of detection, to a pseudocount based on the minimum value observed for that particular feature.

<u>Linear model design</u>

Except where otherwise noted, statistics were performed using linear models implemented with the lme4 package in R and the lmerTest package to generate p-values. For data with repeated measures from the same individual/vessel (human participants, artificial gut), a mixed-effects model was used by including a random intercept term (1 | ID) in the model formula within the lmer () function; otherwise (for *in vitro* cross-reactivity experiments, for which measurements were taken from distinct cultures) the lm () function was used. For the fecal metabolic potential experiment, the data were found to be non-normal by using the descdist () function in the fitdistrplus package; in this case, we therefore used a generalized mixed-effects linear model with a Gamma distribution, implemented with the following call:
fit <- glmer (concentration ~ group*time + (1 | ID), data=hpaec.data, family = Gamma (link = "log"))

For metatranscriptomic data, analysis was done using an ALDEx2 generalized linear model (*57*) with vessel and time as factors. For the within-taxa analysis, an individual GLM was constructed for each taxon in order to account for changes in the total amount of RNA contributed by a given taxon (i.e. to avoid seeing apparent changes in a transcript just because the compositional abundance of that taxon changed). Multiple comparisons correction was performed by the Benjamini-Hochberg method.

The 16S sequencing data from the artificial gut required a more complex model to account for the multivariate and compositional nature of the data as well as the high density of repeated measures samples. We designed an auto-regressive Bayesian multinomial logistic-normal model using a *pibble* model in the *fido* R package (*58*) (Extended Data Fig. 3a). The first order autoregressive AR (1) portion of this model was designed by creating a variable for each sample, and setting the prior covariance of samples taken from the same vessel to $0.8^{\Delta t/8}$, where $\Delta t$ is the number of hours elapsed between the samples. Samples taken from different vessels had a covariance of 0, and all samples had a covariance of 1 with themselves. In addition to these sample variables, we included a $\Delta t$ variable to account for consistent drift over time, and four experimental variables. The values of the four experimental variables supplied in the covariate matrix were all binary (0 or 1), and were only 1 for select values during the Treatment week. The variable $\beta_1$ corresponded to +6 hours from the first dose, $\beta_1$ to +22 hours from the first dose (or, -2 hours from the second dose), $\beta_1$ to +6 hours after the second through fifth doses, and $\beta_1$ to +22 hours from the second through fourth doses (no sample was taken the morning after the fifth dose). All five of these were fixed across vessels, and all had prior covariance of 0 with each other, and 1 with themselves (Extended Data Fig. 5a). Essentially, the variables were so designed in order distinguish the effects at +6 hours and +22 hours, and to uniquely distinguish the effects on Day 1, since we had previously identified unique metabolic activity (or a lack thereof) on this day (Fig. 1b). The model was robust to changes in the priors.

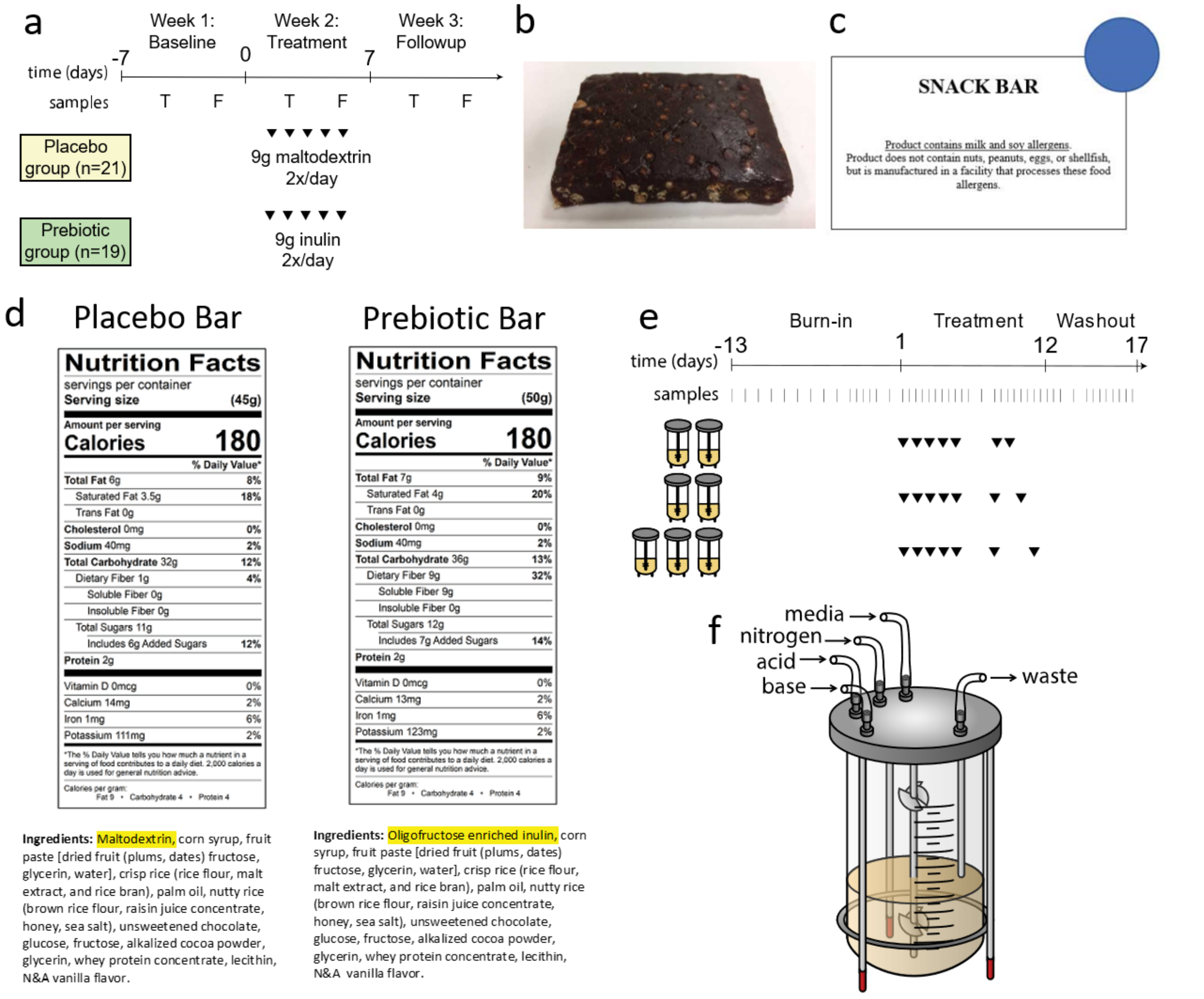


**Fig. S1. Study design.**

**a**, Human participant study design. Participants consumed snack bars twice daily, which contained either 9g inulin (prebiotic group) or 9g maltodextrin (placebo), and provided stool samples twice weekly, on Tuesdays and Fridays. **b**, Photo of snack bar consumed by research participants. **c**, Approximate label used for snack bars. A colored dot, either red or blue, was coded to the identity of the bar, to which participants were blinded. **d**, Nutrition facts and ingredients for the placebo and prebiotic bars. **e**, Schematic of artificial gut run. After a 13-day burn-in period, discrete inulin dosing (denoted by ▼) was started in the seven artificial guts used for the experiment. Samples were taken once or twice daily, denoted by |. **f**, Diagram of the "artificial gut", a continuous-flow bioreactor.

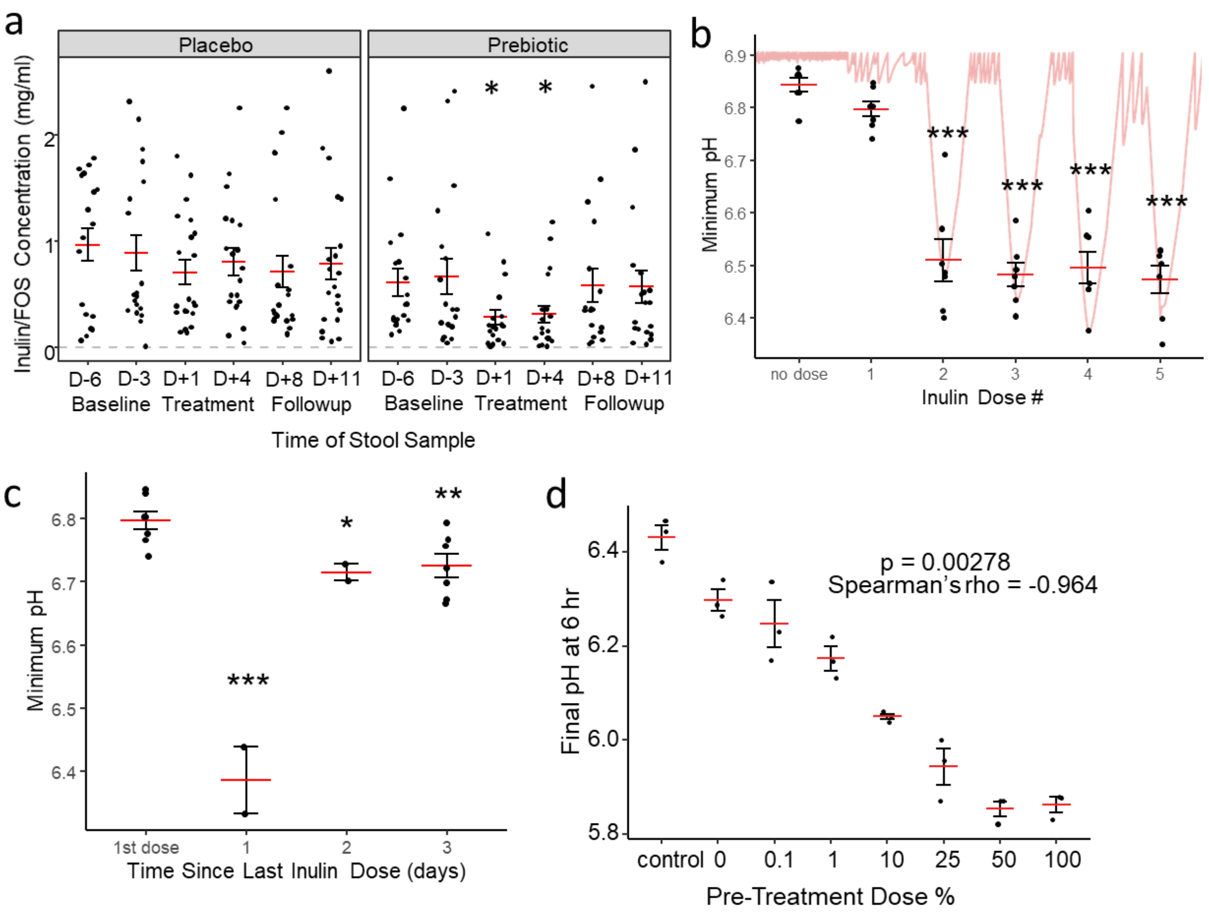


**Fig. S2. Enhancement of inulin metabolism within 24 hours of first dose.**

**a**, Inulin/FOS concentrations (DP 3+) after incubation of slurried stool samples with inulin for 24 hours ex vivo for all time points in both groups. (Prebiotic D+1 p = 0.0282, D+4 p = 0.0191; mixed-effects GLM with Placebo group and D-6 as intercept; Placebo n = 18, 19, 21, 20, 21, 21; Prebiotic n = 19, 19, 18, 19, 18, 19). **b**, Minimum pH reached over the 24 hours following each inulin dose. Continuous pH trace for Vessel 1 shown. (Mixed-effect linear model with "no dose" as intercept; n = 7 vessels.) **c**, Minimum pH reached over the 24 hours following each inulin dose, plotted by time since previous dose. (Mixed effects linear model with 1st dose as intercept; n = 7, 2, 2, and 7 vessels.) **d**, Final pH after 6 hours incubation on inulin, preceded by pre-treatment with an inulin dose of varying concentration. (n = 3 cultures.) Mean and standard error plotted. * $p < 0.05$, ** $p < 0.01$, *** $p < 0.001$.

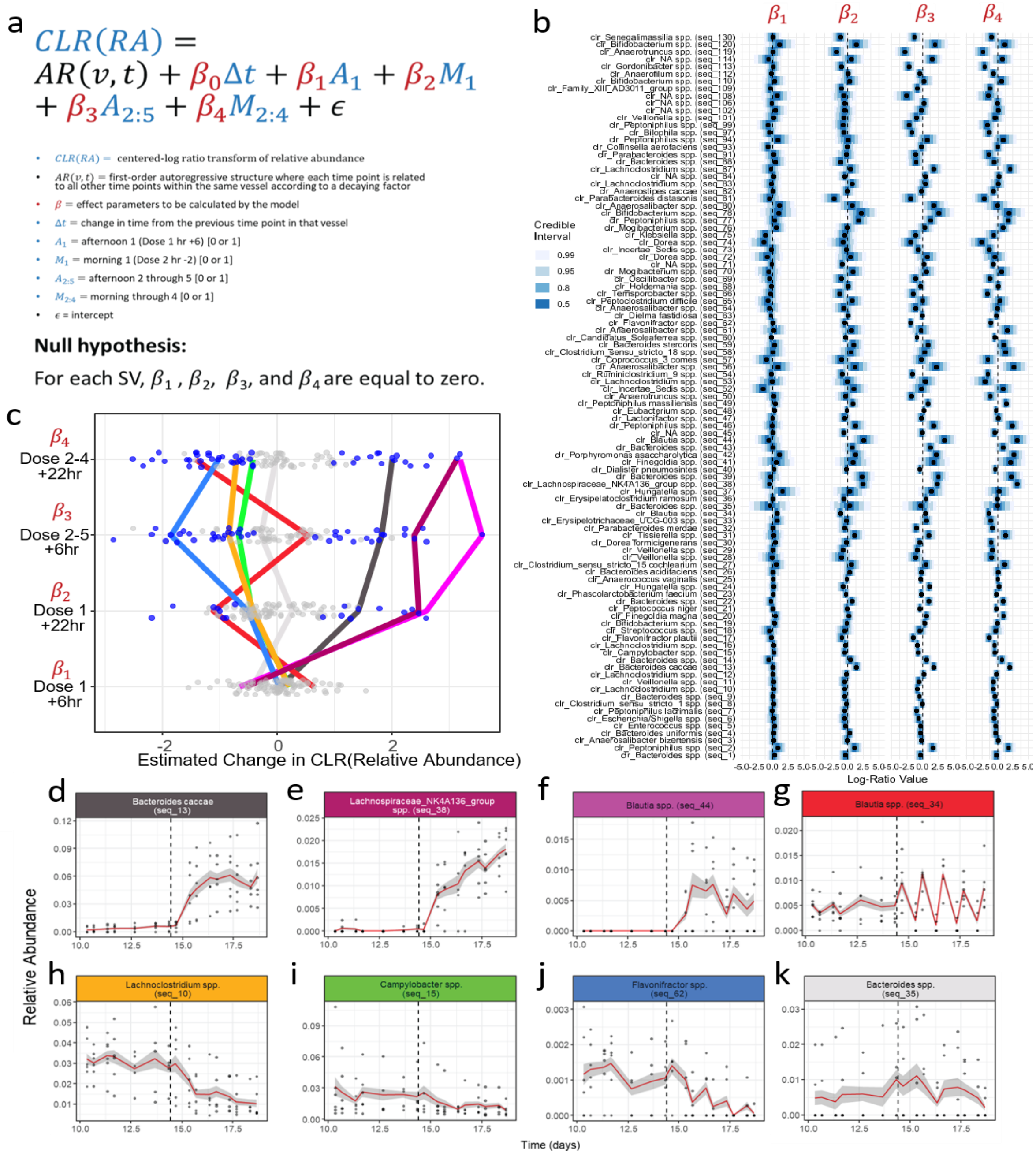


**Fig. S3. Bayesian multinomial logistic-normal model for 16S data.**
**a**, Simplified model equation. Blue terms are supplied to model in the count matrix and covariate matrix, and red terms are calculated by the model. **b**, Parameter estimates for all SVs. SVs are considered significant for a given parameter if the 95% credible interval excludes zero. **c**, Parameter estimates from (**b**), collapsed. Each point is an SV; blue points show SVs for which the 95% confidence interval for that parameter excludes zero. Selected SVs tracked using colored lines. **d-k**, Plots of individual SVs highlighted in (**c**). Examples of taxa significantly increasing with treatment (**d-f**), oscillating (**g**), significantly decreasing with treatment (**h-j**), or not significantly changing (**k**). Vertical lines represent times of first inulin dose. Mean and standard error plotted. (n = 6 vessels.)

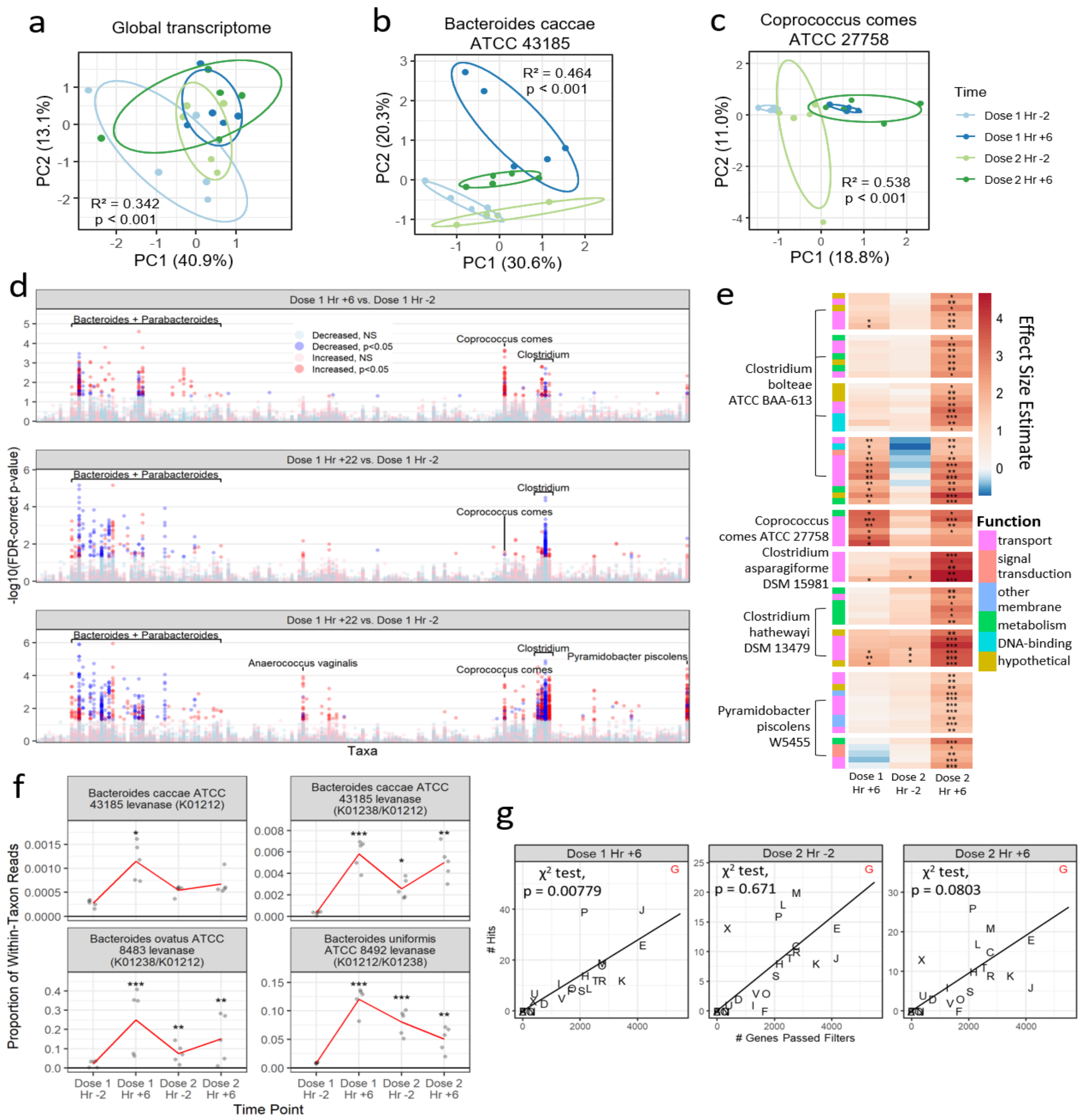


**Fig. S4. Changes in artificial gut metabolome following inulin exposure.**

**a-c**, PCA plots of global transcriptome (**a**) and individual transcriptomes of select taxa, a primary inulin degrader (**b**) and a degrader of secondary metabolites (**c**), by time point. PERMANOVA $R^2$ and p-values shown. **d**, Results of within-taxon ALDEx2 analysis by time point. Taxa clustered along x-axis by phylogeny. **e**, Heatmap of genes comprising putative operons in non-*Bacteroides* species induced on inulin treatment, annotated by general function. Each row is a gene and each cluster is a putative operon. **f**, Gene transcripts significantly increased in the within-taxon analysis and mapping to KEGG categories found to be significantly increased in the global analysis at Dose 1 Hr +6 (K01212 and K01238). Mean shown in red. **g**, Representation of COG categories in transcripts found to be significantly differentially expressed by ALDEx2 GLM at each time point in the within-taxon analysis. The black line describes where all COG categories would lie if they were represented in the hits equally to their proportion in the overall gene set. Chi-squared test for enrichment of category G (carbohydrate metabolism and transport) shown. (n = 5 vessels.) * p < 0.05, ** p < 0.01, *** p < 0.001.

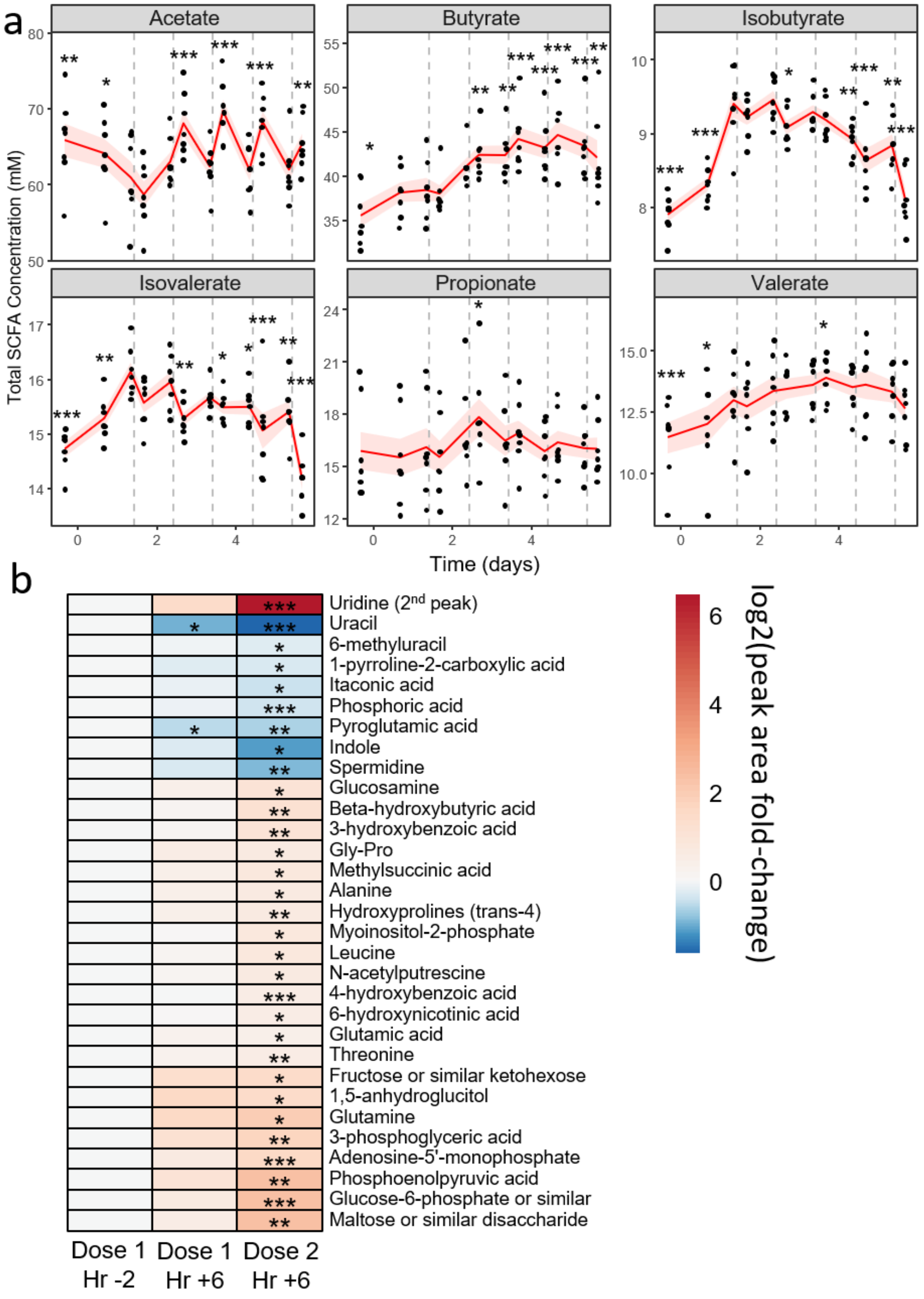


**Fig. S5. Changes in artificial gut metabolome.**

**a**, All measured SCFAs in the artificial gut during dosing week and the two days prior. (Mixed-effects linear model with Dose 1 Hr -2 (Day 1.42 on x-axis) as intercept; n = 7 vessels.) Vertical lines represent times of inulin dosing. Mean and standard error plotted. **b**, GC/MS metabolites found to be significantly different from baseline by mixed-effects linear model in the artificial gut (n = 6 vessels). * $p < 0.05$, ** $p < 0.01$, *** $p < 0.001$.

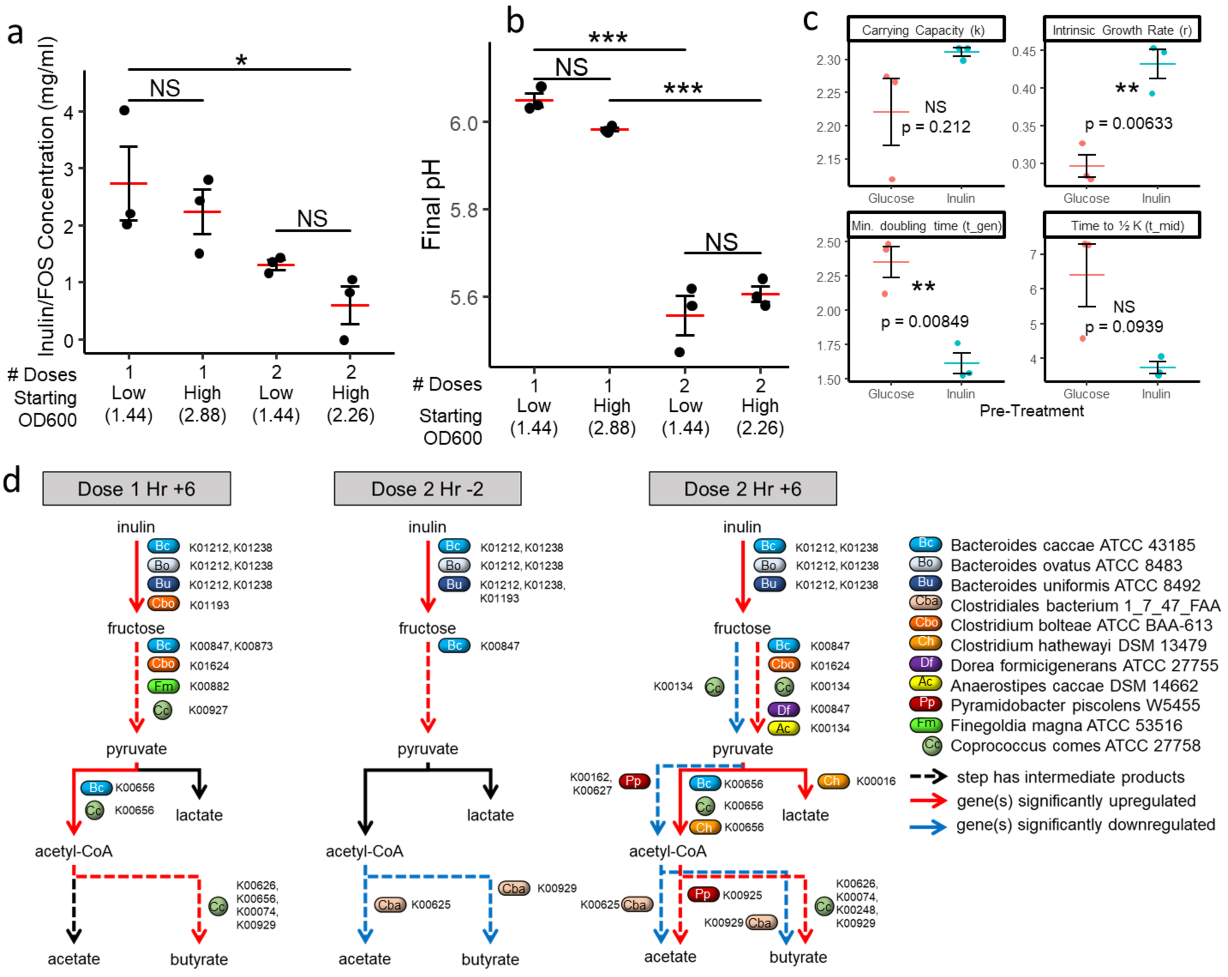

**Fig. S6. Roles of total cell biomass and transcriptional state on ecological memory.**

**a-b**, Effect of starting cell concentration on inulin/FOS breakdown (**a**) and final pH (**b**). (Tukey test; n = 3 cultures.) **c**, Metrics from growth curves shown in Fig. 3c calculated by the growthcurver R package. (Two-sided unpaired t-test; n = 3 cultures.) Mean and standard error shown. **d**, Simplified pathway diagram depicting the conversion of inulin to relevant secondary metabolites by time point. Taxa found to significantly (Benjamini-Hochberg adjusted $p < 0.05$) up- or downregulate genes encoding enzymes carrying out steps in this pathway shown. (ALDEx2 GLM; n = 5 vessels.) NS $p > 0.05$, * $p < 0.05$, ** $p < 0.01$, *** $p < 0.001$.

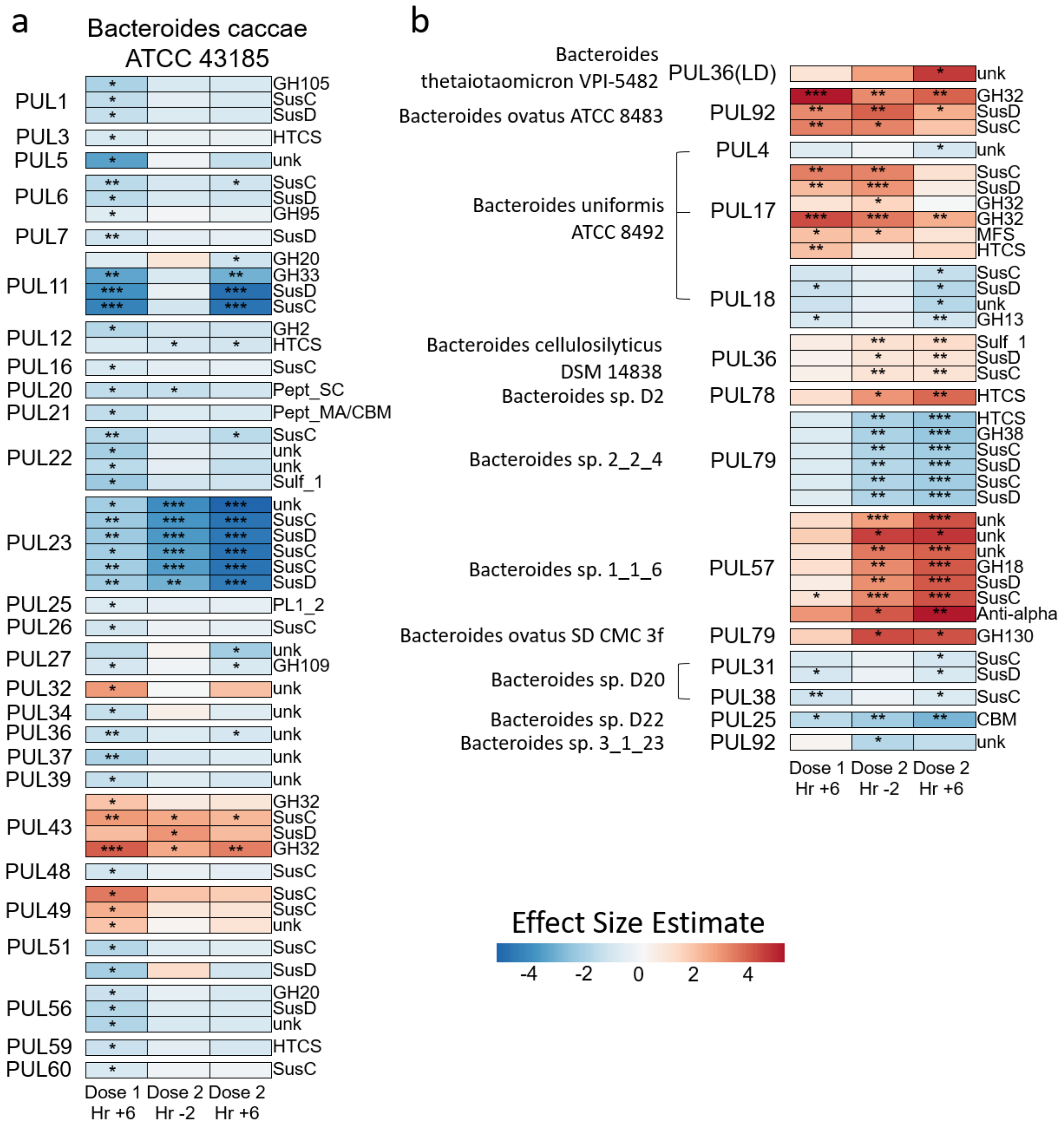


**Fig. S7. Activation of PULs in *Bacteroides*.**

**a-b**, Polysaccharide utilization loci in *Bacteroides caccae* ATCC 43185 (**a**) and all other Bacteroides species (**b**) for which at least one gene was significantly differentially expressed following inulin treatment. Conserved functional prediction from PULDB shown. (ALDEx2 GLM; n = 5 vessels.) * $p < 0.05$, ** $p < 0.01$, *** $p < 0.001$.

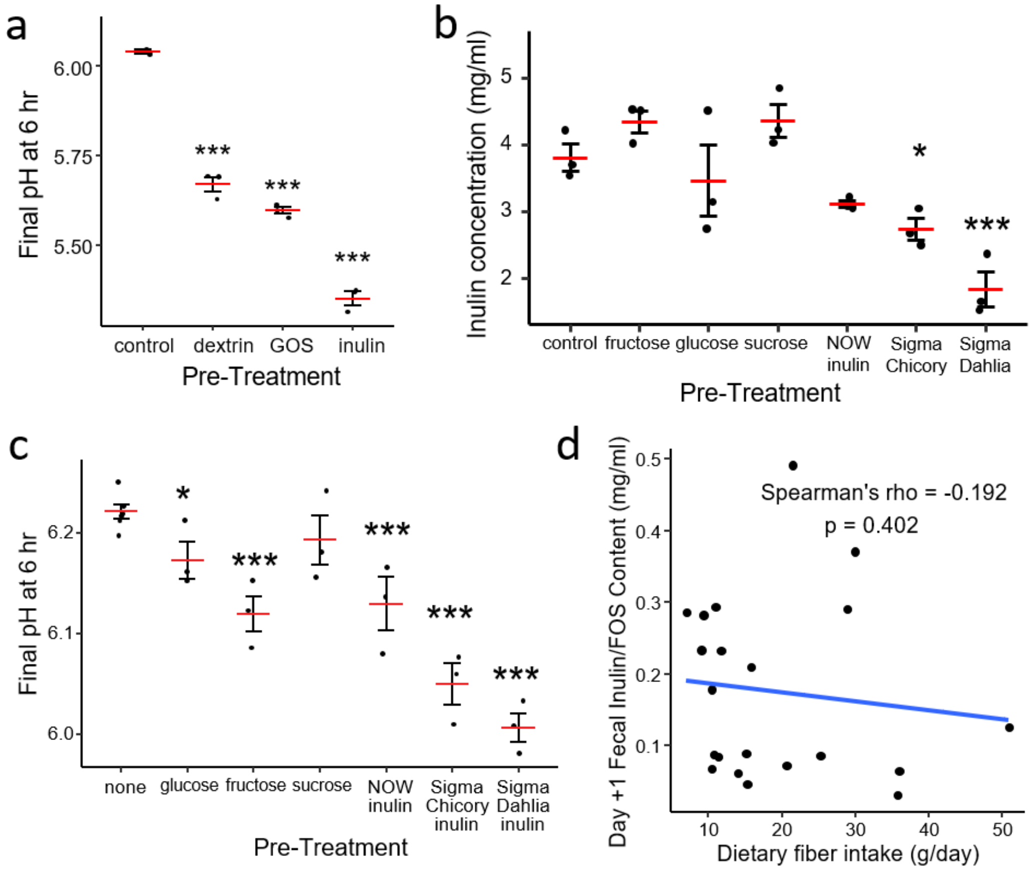


**Fig. S8. Supplementary data on cross-reactivity of ecological memory.**

**a**, Final pH and after inulin treatment and pre-treatment with different prebiotics. (Linear model with control as intercept; n = 3 cultures.) **b-c**, Final inulin concentration (**b**) and pH (**c**) when pre-treated with inulin of various chain length or component mono/di-saccharides. (Linear models with control as intercept; n = 3 cultures.) **d**, Correlation between participant baseline dietary fiber (DHQ3) intake and fecal inulin/FOS content on Treatment Day +1 in the Placebo group (n = 21 participants). * $p < 0.05$, ** $p < 0.01$, *** $p < 0.001$.

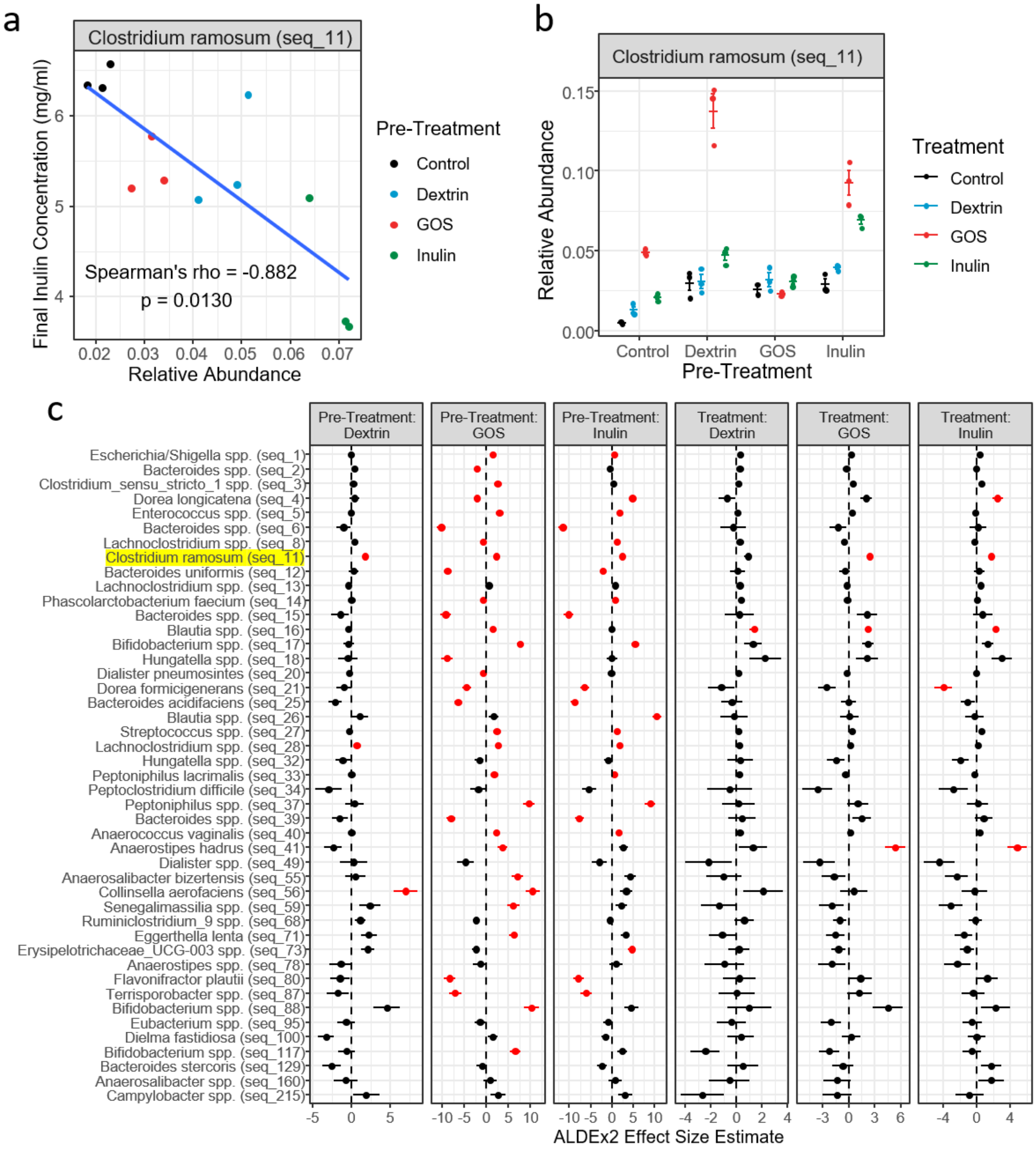


**Fig. S9. 16S sequencing results for prebiotic crossover experiment.**
**a**, Correlation of *Clostridium ramosum* relative abundance with final inulin concentration. This was the only SV for which a significant correlation was found by ALDEx2 test of Spearman correlation (after Benjamini-Hochberg correction). **b**, Relative abundance of *C. ramosum* by pre-treatment and final treatment. Mean and standard error shown. **c**, ALDEx2 GLM results for all SVs. Mean effect size estimate and standard error shown. Red color indicates a statistically significant ($p < 0.05$) result for a given SV-covariate combination. Note that *C. ramosum* (along with one other SV mapping to *Lachnoclostridium*) has positive and significant values for pre-treatment with all three prebiotics. (n = 3 cultures.)

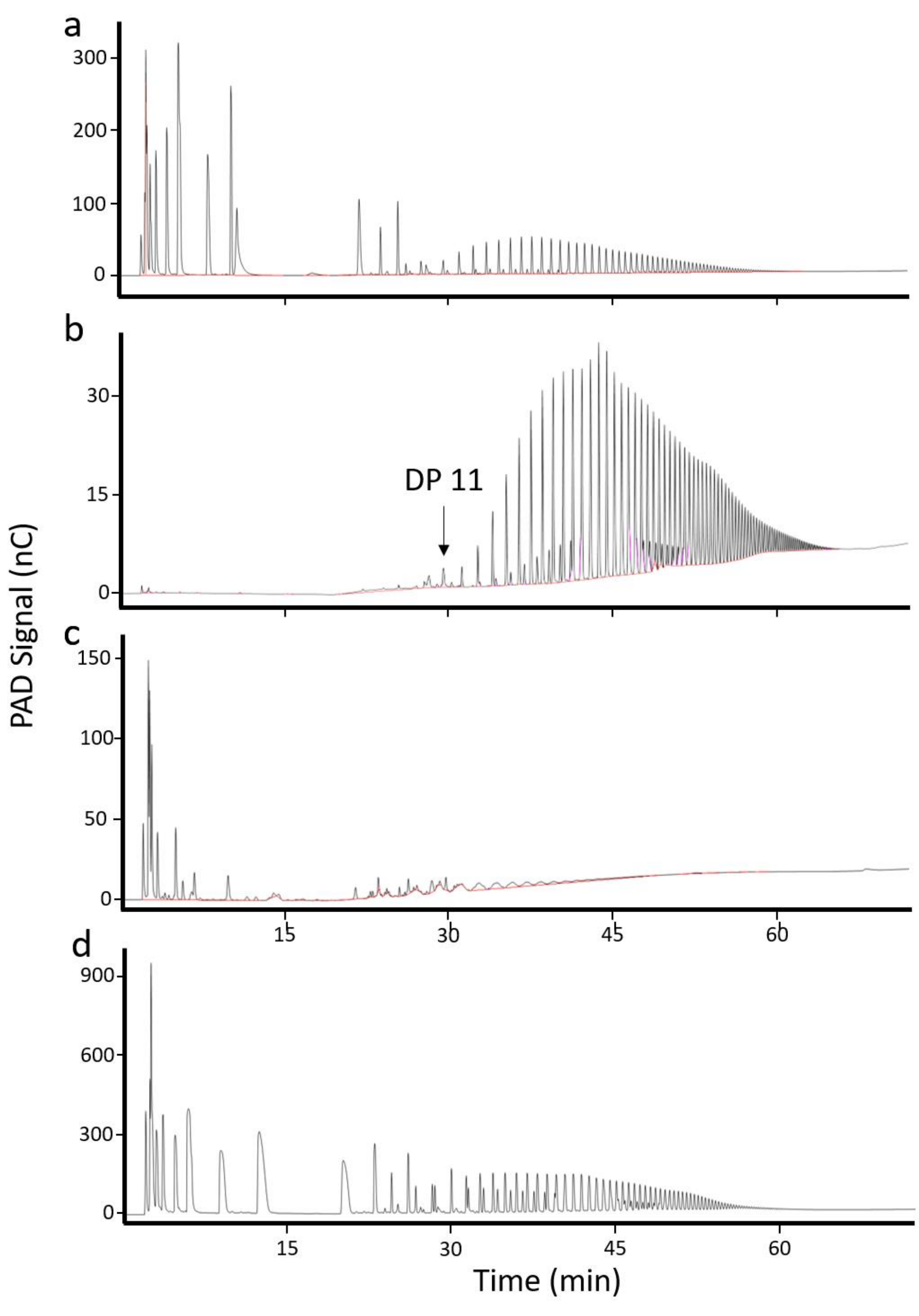


**Fig. S10. Spectra of primary inulin sources used in this study.**

**a,** Orafti Synergy1 inulin, a mixture of short- and long-chain inulin, used for the human study. **b**, Sigma Inulin from dahlia tubers, a very pure source of long-chain inulins, used for most *in vitro* work including the artificial gut experiment. **c**, NOW inulin, a source with mostly shorter chain FOS. **d**, Jarrow inulin, another mixed chain length inulin.
